# Riboflavin-sensitized UVA collagen crosslinking produced a controllable, dose-dependent increase in the nanomechanical strength of ex vivo bovine dura mater

**DOI:** 10.64898/2026.06.03.729951

**Authors:** Ioannis Vasilikos, Sukesh Mysore Swamy, Ulrich G. Hofmann, Ulrich Hubbe, Roland Rölz, Angeliki Stathi, Katerina Wolk, Daniel Strahnen, Amir El Rahal, Mukesch J. Shah, Jürgen Grauvogel, Florian Volz, Boris Mizaikoff, Lorena Diaz, Vidhya M. Ravi, Kevin Joseph, Jürgen Beck

## Abstract

**Background:** Dural defects, either from trauma, tumor resection, surgical approaches, genetics, or spontaneously represent a significant clinical challenge in neurosurgery. Given the established efficacy of riboflavin-sensitized ultraviolet-A (UVA) photo-crosslinking in ophthalmology, this study investigated its feasibility and dose-response characteristics as a novel strategy to biochemically augment the mechanical integrity and strength of ex-vivo bovine dura mater.

**Methods:** Forty bovine dura mater specimens were treated ex vivo using riboflavin concentrations of 2, 4, or 8 mM combined with UVA irradiation at 0.3 or 3 mW/cm². PBS-treated specimens exposed to UVA served as controls. Atomic force microscopy nanoindentation was used to measure the local elastic modulus in matched regions before and after treatment, enabling paired assessment of treatment-induced mechanical changes while minimizing inter-sample variability. Post-treatment stiffness, fold-change from baseline, and riboflavin dose-response relationships were analyzed statistically.

**Results:** Baseline elastic moduli were equivalent across all groups (mean approximately 52 kPa, p=0.92). While UVA alone caused a modest approximately 2- to 3-fold stiffness increase, riboflavin-UVA treatment produced a dramatic, concentration-dependent effect. The highest treatment (8 mM RF, 3 mW/cm² UVA) increased the elastic modulus 150-fold, from approximately 53 kPa to approximately 8,000 kPa. Post-UV stiffness exhibited a strong linear relationship with riboflavin concentration (R² = 0.994), indicating a precisely titratable crosslinking effect. All treatment conditions were statistically distinguishable (p < 0.001).

**Conclusion:** Riboflavin-sensitized UVA crosslinking substantially increases the nanomechanical strength of ex vivo bovine dura mater in a controllable, dose-dependent manner. These findings establish a proof of concept for biochemical reinforcement of dural tissue that might be used clinically. As a next step evaluation using human dura, macroscopic biomechanical testing, penetration-depth analysis, and safety assessment is warranted.

## Introduction

The dura mater is a collagen-rich membrane that provides structural protection to the central nervous system and contributes to the maintenance of a watertight cerebrospinal fluid compartment. In neurosurgery, integrity may be compromised by trauma, tumor resection, spontaneous intracranial hypotension (SIH) or the surgical approach itself represents a significant clinical challenge. Failure to achieve a durable, watertight dural closure can lead to severe complications, including CSF leakage, meningitis, pseudomeningocele formation, and herniation of spinal rootlets, spinal cord and even the cerebrum[1]. These risks are particularly elevated in specific clinical scenarios, such as in patients with connective tissue disorders like Marfan syndrome that cause inherent dural weakness (dural ectasia), in revision surgeries where the dura is scarred and friable, and following the resection of dural-invasive tumors that necessitate complex reconstructions[2]. A method to biochemically enhance the strength of the dura mater would therefore be of substantial clinical value.

The ultrastructure of the dura mater is characterized by a dense, multi-layered connective tissue architecture composed primarily of type I collagen, with smaller amounts of type III collagen contributing to tissue elasticity[2–4]. Electron microscopy reveals an outer fibroelastic layer, a middle fibrous layer rich in collagen, and an innermost border cell layer with minimal extracellular collagen[3] The collagen fibers are predominantly oriented longitudinally and consist of large, non-uniform fibrils similar to those found in tendons, conferring anisotropic mechanical properties with greater tensile strength in the longitudinal direction[5–7].

A well-established method for strengthening collagenous tissue is riboflavin-sensitized ultraviolet-A (UVA) light-induced collagen cross-linking. This technique has become the standard of care in ophthalmology for treating keratoconus, a condition where a weakened cornea progressively deforms[8–10]. The combination of riboflavin and UVA light creates new covalent bonds within the stromal collagen, increasing the tissue’s biomechanical stiffness and halting disease progression[9,11,12]. We have previously demonstrated that this principle is transferable from the avascular cornea to the intervertebral disc, another collagen-rich tissue, showing that riboflavin-UVA treatment can significantly augment the mechanical properties of the disc’s extracellular matrix [13].

For this initial feasibility study, atomic force microscopy nanoindentation was selected to quantify local nanomechanical changes after crosslinking. AFM enables high-resolution assessment of tissue surface mechanics and has previously been applied to characterize structural and mechanical heterogeneity in dura mater [14,15,16,17,18]. Although AFM does not measure bulk tensile strength or functional repair performance, it provides a sensitive method to determine whether riboflavin-UVA treatment can alter local dural stiffness under controlled ex vivo conditions.

In this study, we investigated whether riboflavin-sensitized UVA crosslinking increases the nanomechanical stiffness of ex vivo bovine dura mater. Using paired AFM measurements before and after treatment, we evaluated the effects of different riboflavin concentrations and UVA power densities. We hypothesized that riboflavin-UVA exposure would produce a dose-dependent increase in local dural stiffness, thereby establishing an initial proof of concept for photochemical modulation of dural tissue mechanics.

## Materials and Methods

### Ethics statement

This article does not contain any studies with human participants performed by any of the authors. All applicable international, national, and institutional guidelines for the care and use of animals were followed. Our in vitro experiments were conducted in a manner that did not violate any provisions of the ARRIVE guidelines or the Declaration of Helsinki regarding animal studies. The bovine dura material used in the study was obtained from commercial meat processing, which conforms to the European regulatory standards. Following consultation with the University of Freiburg’s local ethics committee, it was determined that formal approval of the study was not required (26-1235).

### Tissue source and sample preparation

40 bovine dura preparations (5 cm × 5 cm) were harvested from 4- to 6-month-old bovine skulls, from the central meat processing unit in the city of Freiburg and kept refrigerated at 4 °C in saline. All experiments were conducted within 48hrs post sample acquisition.

We chose to investigate the RF concentrations and UVA intensities that we have previously investigated in our work with the bovine intervertebral discs [13].

Therefore, we grouped the samples as follows:

Group A (n=5): 2 mM RF (25 measurements) and UVA 0.3 mW/cm2 (25 measurements)

Group B(n=5): 2 mM RF (25 measurements) and UVA 3 mW/cm2 (25 measurements)

Group C(n=5): 4 mM RF (25 measurements) and UVA 0.3 mW/cm2 (25 measurements)

Group D(n=5): 4 mM RF (25 measurements) and UVA 3 mW/cm2 (25 measurements)

Group E(n=5): 8 mM RF (25 measurements) and UVA 0.3 mW/cm2 (25 measurements)

Group F(n=5): 8 mM RF (25 measurements) and UVA 3 mW/cm2 (25 measurements)

Control Group I (n=5): Dura Sample in PBS solution and UVA 0.3 mW/cm2 (25 measurements)

Control Group II (n=5): Dura Sample in PBS solution and UVA 3 mW/cm2 (25 measurements)

### Riboflavin exposure

Grouped dura specimens were submerged into PBS (Control I and II) or 2 mM, 4mM and 8 mM RF solution respectively (groups A-F) for 15 min in a dark environment (to avoid unintended RF activation from natural light sources) and then placed flat on a petri-dish.

### UVA irradiation

UVA irradiation was performed using an UVA-LED module (Opsytec Dr. Gröbel GmbH, Ettlingen, Germany). The UVA-LED module produced a homogenous field of UVA light at a wavelength of 365–370 nm. UVA-light intensity was adjusted by placing the UVA-LED module at different distances from the cell culture plate, based on preparatory calibrating experiments. UVA-absorbance of the petri-dish plates was also considered. This approach enabled dura mater specimen irradiations with 3 mW/cm2 for 15 min. The intensity was calibrated using a radiometer (RM-12, Opsytec Dr. Gröbel).

### Atomic Force microscopy

To probe the nanoscale mechanics of dura mater and determine how riboflavin-mediated crosslinking alters its behavior, we employed atomic force microscopy (AFM) [14]. In this type of ultramicroscopy, a finely pointed probe indents the tissue surface while a laser-photodiode system precisely tracks cantilever deflection, yielding force-distance curves that reveal the material’s elastic modulus and highlight subtle reorganizations in the collagen network after treatment [15,16]. All measurements and high-resolution mechanical maps were acquired on a Bruker Bioscope Resolve 9.0 instrument, using both PeakForce Quantitative Nanomechanical Mapping (PF-QNM) and conventional ramping modes to combine detailed imaging with quantitative indentation data [17,18]. Triplicate sets of data were collected from each measured region.

The Bioscope AFM is integrated with an inverted optical microscope for precise positioning and sample visualization [5]. Bio MLCT probes (Bruker Corporation) were used throughout the study [6]. The spring constant of each cantilever was calibrated using the thermal tune method prior to measurements [7]. Bio MLCT probes have a nominal spring constant range of 0.01–0.06 N/m and a tip radius of 20–60 nm, making them suitable for soft biological tissue measurements [6].

Indentation forces were maintained within the range of 5 nN to 40 nN to avoid damaging the sample [19]. Despite the relatively stiff nature of dura mater, indentation depths were carefully limited to less than 10% of the sample thickness to minimize substrate influence and maintain the validity of the contact mechanics models [15]. PF-QNM mode was operated at a frequency of 1 kHz, allowing simultaneous acquisition of topographical and nanomechanical data. Multiple force distance curves were acquired per region to ensure statistical robustness. Samples were immersed in phosphate-buffered saline (PBS) during AFM measurements to preserve physiological conditions. Samples were mounted on an inverted microscope stage and manually positioned in the x and y directions to target regions of interest. To evaluate mechanical changes due to crosslinking, identical regions were measured before and after riboflavin treatment. This minimized inter-sample variability and allowed direct comparison of mechanical behavior before and after crosslinking.

Force-distance curves were analyzed using contact mechanics models appropriate for the probe geometry. Given that Bio MLCT probes feature pyramidal tip geometry, the Sneddon contact model was applied for elastic modulus extraction[20]. Only high-quality curves exhibiting clearly defined contact points, smooth approach curves, and strong correlation between experimental data and theoretical model (R2 > 0.95) were included in modulus calculations. Curves showing evidence of adhesion, surface contamination, or poor contact were systematically excluded from analysis to ensure measurement reliability.

The sequence for AFM preparation and calibration is illustrated in **Fig. 1**, and the experimental workflow utilized is outlined in **Fig. 2**.

**Fig. 1.**
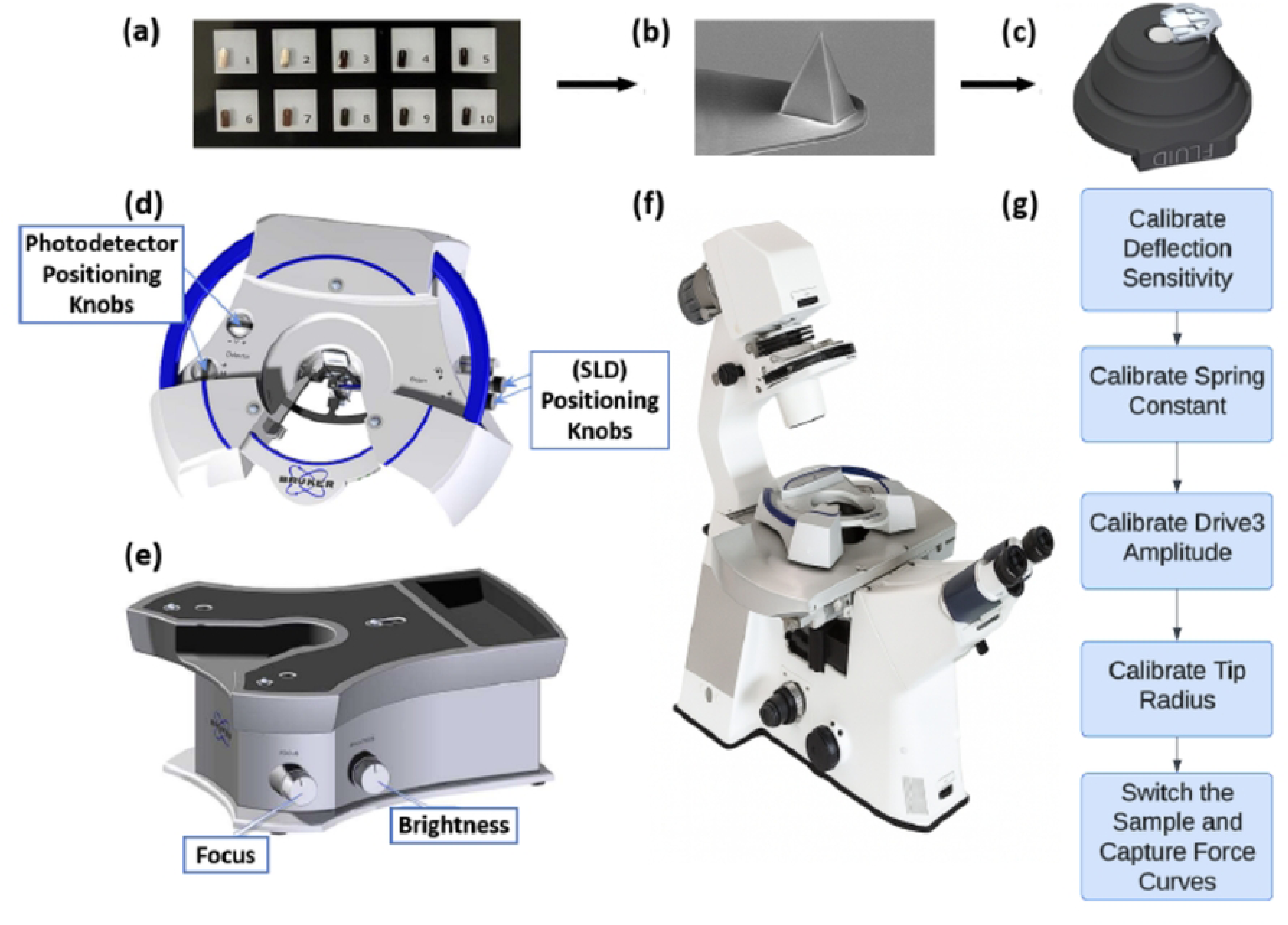
AFM experimental workflow: (a) Box containing custom manufactured Bruker cantilever, (b) Image of an isolated cantilever showing the probe, (c) Probe holder onto which the cantilever is loaded (d) AFM head (shown with laser alignment features) onto which the probe holder is mounted, (e) Base supporting laser alignment, (f) optical microscope featuring the AFM head, and (g) calibration sequence including deflection sensitivity, spring constant, drive amplitude, and tip radius measurements before force curve acquisition. Panels (b-f) adapted from [6].

**Fig. 2.**
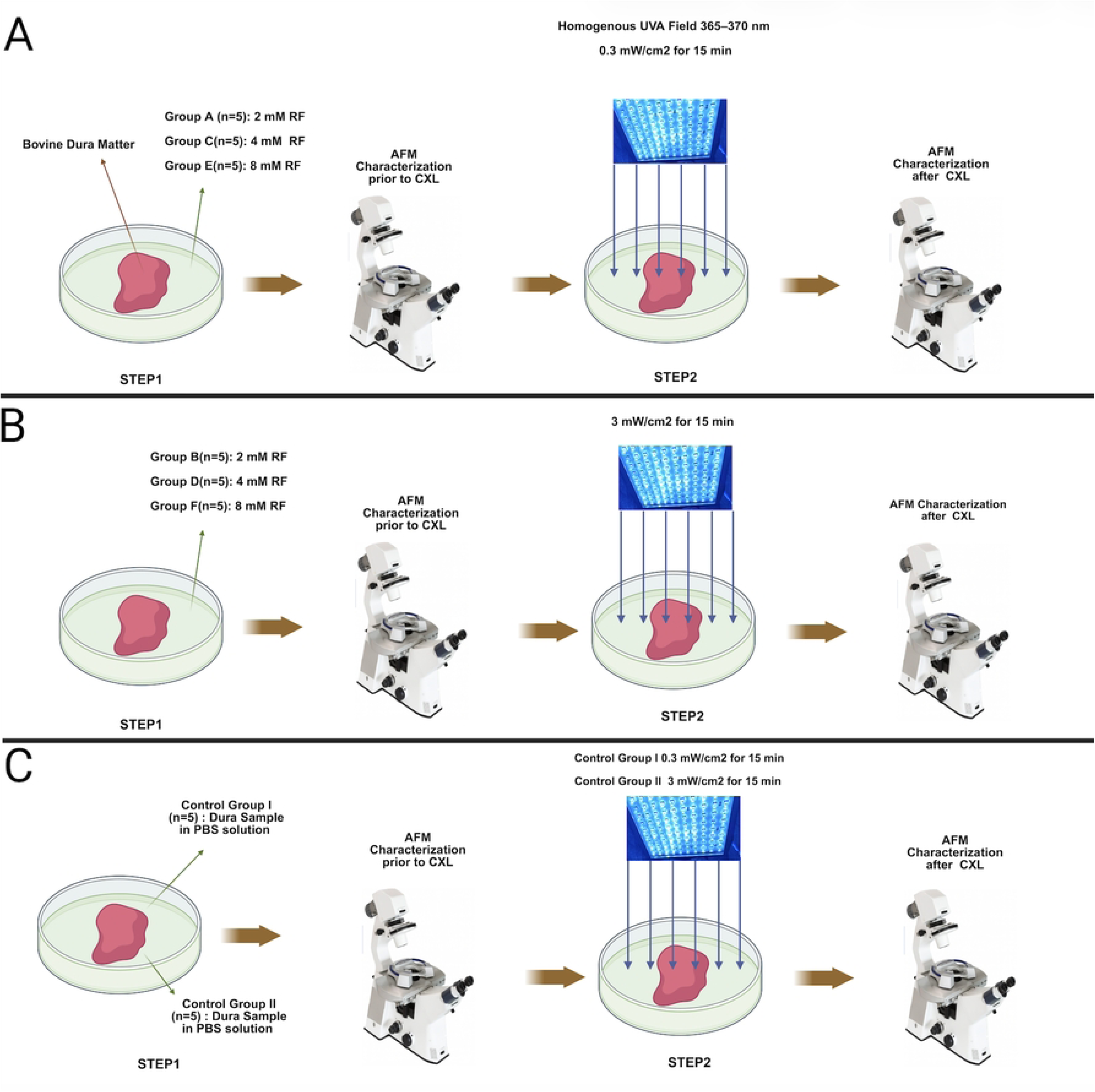
Schematic overview of the experimental workflow: (A) Samples assigned to Groups A, C, and E were treated with riboflavin (RF) at concentrations of 2, 4, and 8 mM, respectively. Following RF application, baseline nanomechanical properties were assessed by atomic force microscopy (AFM). Samples were subsequently exposed to a homogeneous UVA field (365–370 nm, 0.3 mW/cm²) for 15 minutes, after which AFM measurements were repeated to evaluate post-crosslinking changes. (B) Samples assigned to Groups B, D, and F were treated with riboflavin (RF) at concentrations of 2, 4, and 8 mM, respectively. Following RF application, baseline nanomechanical properties were assessed by atomic force microscopy (AFM). Samples were subsequently exposed to a homogeneous UVA field (365–370 nm, 3 mW/cm²) for 15 minutes, after which AFM measurements were repeated to evaluate post-crosslinking changes. (C) Control group samples I and II received Phosphate-Buffered Saline (PBS) only, followed by baseline assessment of nanomechanical properties by AFM. Samples from control group I were subsequently exposed to a homogeneous UVA field (365–370 nm, 0.3 mW/cm²) for 15 minutes, while those from control group II received an equivalent exposure at a higher irradiance (365–370 nm, 3 mW/cm²) for 15 minutes. Post-irradiation AFM measurements were then repeated across all samples in both control groups to evaluate irradiation-induced changes in nanomechanical properties. Created in BioRender. Joseph, K. (2026) https://BioRender.com/8q965kq

### Statistical analysis

All values were converted to kPa for analysis. Fold-change was calculated for each individual measurement as the ratio of post-UV stiffness to baseline stiffness. All statistical analyses were performed using Python (version 3.9.6) with the following packages: pandas (v1.5.3) for data manipulation, NumPy (v1.26.4) for numerical computations, SciPy (v1.13.1) for statistical testing, and statsmodels (v0.14.6) for multiple comparison procedures. Figures were generated using matplotlib (v3.9.4) and seaborn (v0.13.2). Continuous variables are reported as mean +/− standard error of the mean (SEM) for group comparisons and graphical display and mean +/− standard deviation (SD) for characterizing data distributions. Exact p-values are reported; for very small values, scientific notation is used. Statistical significance was defined as p < 0.05. All code and data are available in the project repository (https://github.com/kevinj24fr/dura-photocrosslinking).

### Baseline Equivalence

To verify that treatment groups had equivalent mechanical properties prior to UV exposure, baseline elastic modulus values were compared across all four conditions using one-way analysis of variance (ANOVA). A non-significant result (p > 0.05) was interpreted as evidence supporting baseline equivalence, validating subsequent between-group comparisons of treatment effects.

### Within-Group Treatment Effects

To assess whether UV exposure significantly altered tissue stiffness within each treatment condition, paired (dependent) two-tailed t-tests were performed comparing baseline measurements to post-3 mW/cm² UV measurements for each sample. The paired design accounts for inter-sample variability by using each sample as its own control. Mean differences with 95% confidence intervals (CI) were calculated for each condition.

### Between-Group Comparisons

Post-UV stiffness values were compared across all treatment conditions using one-way ANOVA followed by Tukey’s Honest Significant Difference (HSD) test for pairwise comparisons, which controls the family-wise error rate (FWER) at alpha = 0.05. Homogeneity of variance was assessed using Levene’s test. Because post-UV variance differed significantly across groups (Levene’s W = 19.40, p < 0.0001), Welch’s t-tests were additionally performed as a sensitivity analysis to confirm that conclusions were robust to heteroscedasticity.

### Dose-Response Analysis

The relationship between riboflavin concentration and post-UV tissue stiffness was modeled using ordinary least squares (OLS) linear regression on individual measurements from riboflavin-treated samples (excluding PBS control). Model fit was assessed using the coefficient of determination (R^2), and the slope was tested against the null hypothesis of zero using an F-test. The regression was also performed on group means to assess linearity at the condition level.

### Effect Size Estimation

To quantify the practical significance of treatment effects independent of sample size, Cohen’s d was calculated for each riboflavin condition relative to PBS control using pooled standard deviations. Ninety-five percent confidence intervals for Cohen’s d were computed using the standard error approximation. Effect sizes were interpreted using conventional thresholds: |d| < 0.2 negligible, 0.2-0.5 small, 0.5-0.8 medium, and > 0.8 large.

## Results

### Baseline nanomechanical properties were comparable across treatment groups

A total of 350 AFM measurements were obtained across four treatment conditions: PBS control (n = 50), 2 mM riboflavin (n = 25), 4 mM riboflavin (n = 25), and 8 mM riboflavin (n = 25), (**Fig. 3**). Prior to UV exposure, all groups exhibited comparable elastic moduli (PBS: 51.2 ± 1.9 kPa; 2 mM RF: 52.7 ± 2.8 kPa; 4 mM RF: 52.3 ± 2.8 kPa; 8 mM RF: 53.4 ± 2.7 kPa; values are mean ± SEM). One-way ANOVA confirmed no significant difference in baseline stiffness across groups (F(3, 121) = 0.17, p = 0.92; **Supplementary Fig. 1A**), with group standard deviations ranging from 13.1 to 13.8 kPa (coefficient of variation ≈26%).

**Fig. 3.**
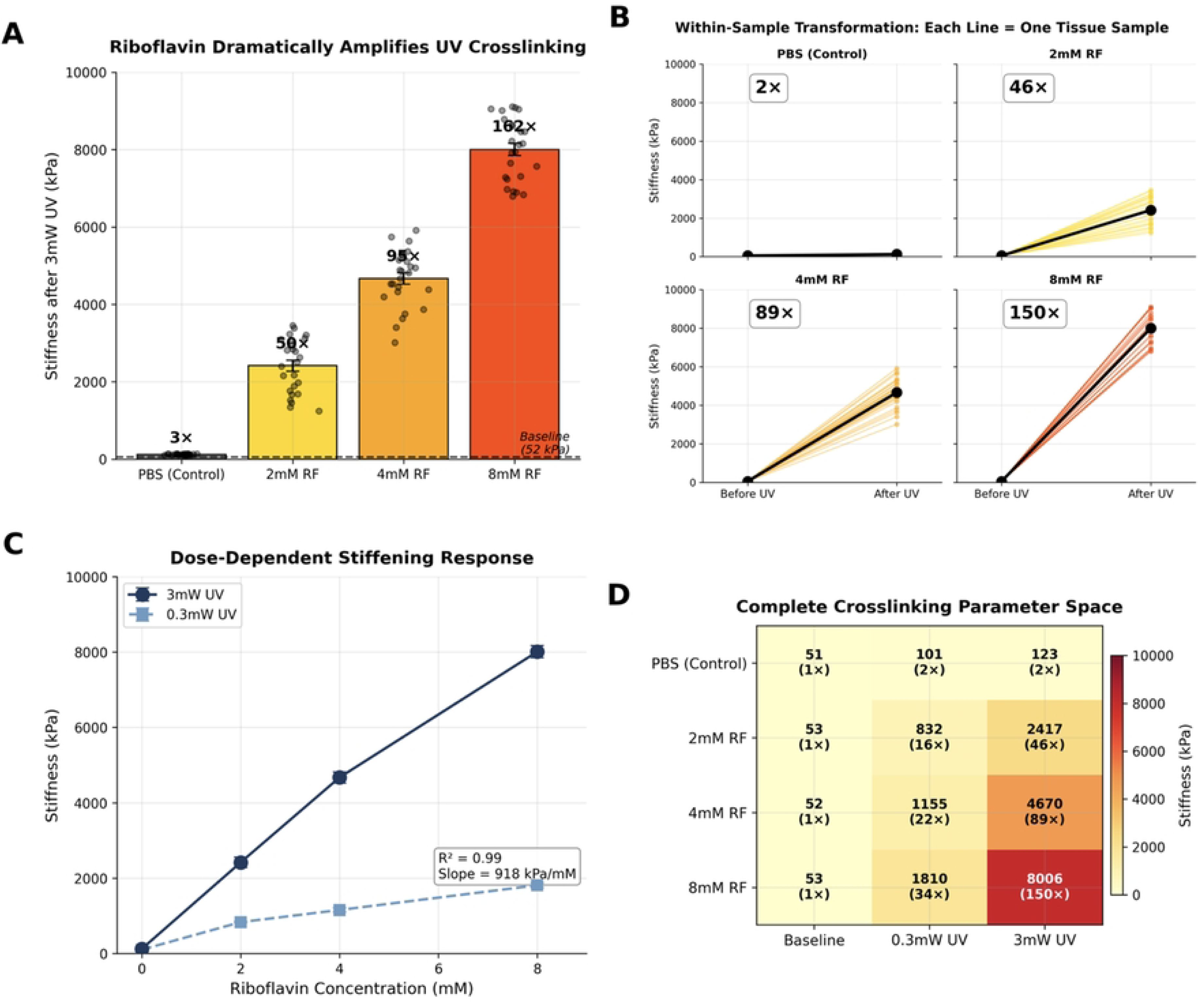
Riboflavin associated and dose dependent nanomechanical strength increase Riboflavin associated nanomechanical strength increase: (A) Elastic modulus (Young’s modulus) of dura mater tissue after 3 mW/cm² UV-A exposure across four treatment conditions. Bars represent mean +/− SEM; individual data points are overlaid (black circles). Fold-change relative to mean baseline stiffness (∼52 kPa, dashed line) is indicated above each bar. PBS control (gray) shows minimal UV-induced stiffening (3x), whereas riboflavin-treated samples exhibit dose-dependent increases of 50x (2 mM, gold), 95x (4 mM, orange), and 162x (8 mM, red orange). n = 25 measurements per riboflavin condition; n = 50 for PBS control. All pairwise comparisons between conditions were significant (Tukey HSD, adjusted p < 0.001). (B) Paired before-and-after measurements demonstrating within-sample stiffness transformation. Each thin colored line connects the baseline and post-3 mW/cm² UV measurement for a single tissue sample; bold black lines with filled circles indicate group means. Fold-change of the group mean is shown in the inset of each panel. All four conditions share a common y-axis (0-10,000 kPa) to enable direct visual comparison. PBS control samples show nearly flat trajectories (2x), while riboflavin-treated samples show steep, uniform increases (46-150x), confirming that stiffening occurs within individual tissue specimens with no overlap between pre- and post-UV distributions. Dose dependent nanomechanical strength increase: (C) Dose-response relationship between riboflavin concentration and post-UV tissue stiffness at two UV power densities. Dark navy circles with solid line: 3 mW/cm² UV; light blue squares with dashed line: 0.3 mW/cm² UV. Error bars represent SEM. At 3 mW/cm², stiffness increases almost linearly with riboflavin concentration (R^2 = 0.99 on group means; slope = 918 kPa/mM), demonstrating predictable, titratable crosslinking. Higher UV power produces proportionally greater stiffening at all riboflavin concentrations. (D) Heatmap summarizing the complete crosslinking parameter space. Rows represent treatment conditions (PBS control through 8 mM riboflavin); columns represent measurement time points (baseline, post-0.3 mW/cm² UV, post-3 mW/cm² UV). Each cell displays the group mean stiffness in kPa with fold-change relative to baseline in parentheses. Color intensity (yellow-orange-red scale) corresponds to stiffness magnitude (0-10,000 kPa). The progressive color gradient across both axes illustrates the interaction between riboflavin concentration and UV power, with maximum stiffening (8,006 kPa, 150x) at 8 mM riboflavin with 3 mW/cm² UV.

**Supplementary Fig. 1.**
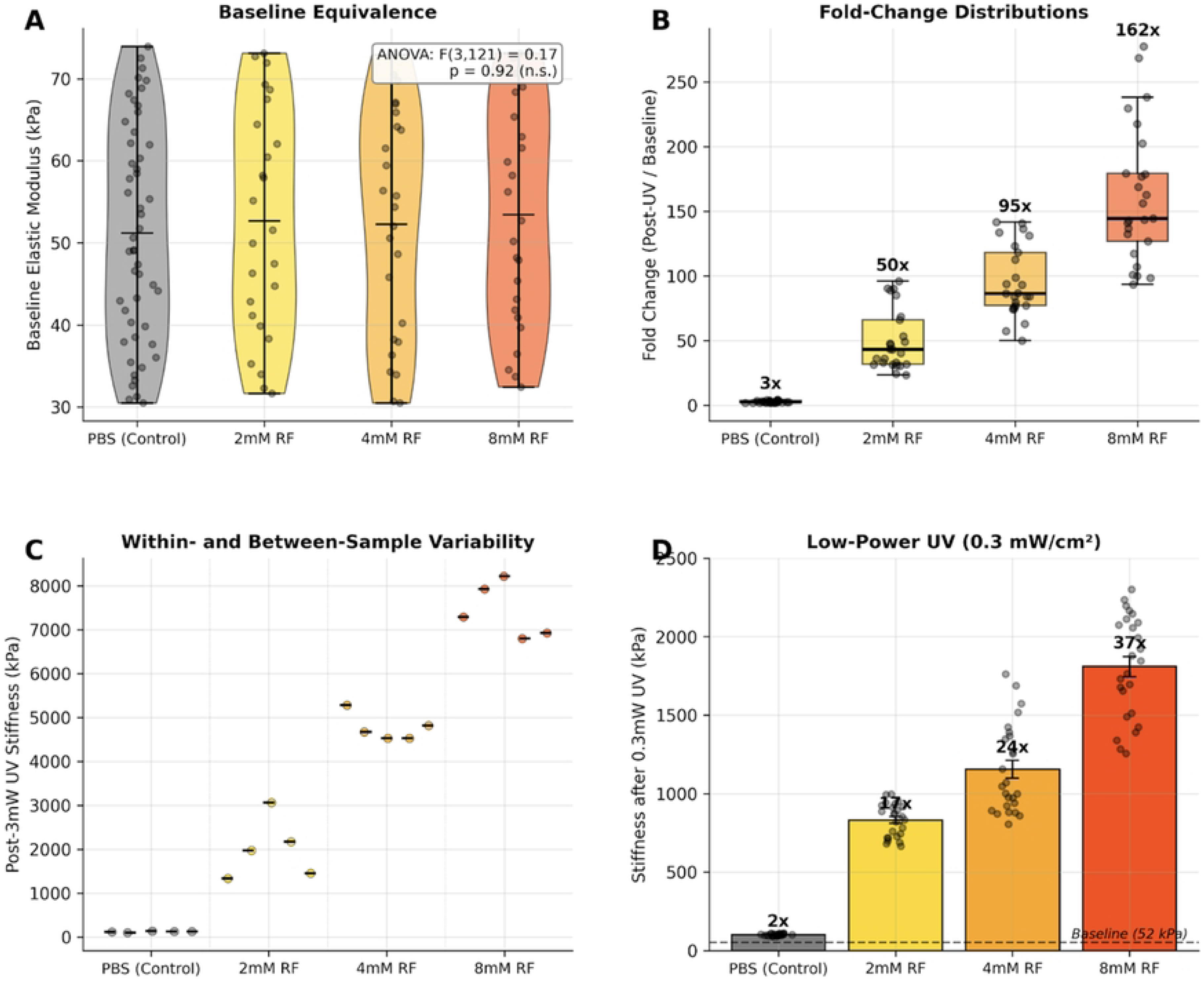
Validation of experimental design, effect magnitude distributions, measurement reproducibility, and low-power UV crosslinking. (A) Violin plots of baseline elastic modulus across treatment groups prior to UV exposure. Horizontal lines indicate group means; individual measurements are overlaid. Distributions are nearly identical (PBS: 51.2 +/− 1.9 kPa, n = 50; 2 mM RF: 52.7 +/− 2.8 kPa, n = 25; 4 mM RF: 52.3 +/− 2.8 kPa, n = 25; 8 mM RF: 53.4 +/− 2.7 kPa, n = 25; values are mean +/− SEM), confirmed by one-way ANOVA (F(3, 121) = 0.17, p = 0.92, n.s.). This validates that post-treatment differences are attributable to photo-crosslinking rather than pre-existing tissue variability. (B) Box-and-whisker plots of individual fold-change values (post-3 mW/cm^2^ UV stiffness divided by baseline) with overlaid data points. Boxes span the interquartile range; horizontal lines indicate medians; whiskers extend to 1.5x IQR. Mean fold-change is annotated above each group. PBS control shows a narrow distribution centered at 3x, while riboflavin-treated conditions show progressively larger fold-changes (50x, 95x, 162x) with increasing spread at higher concentrations, reflecting the greater dynamic range of crosslinking. (C) Nested dot plot showing post-3 mW/cm^2^ UV stiffness for each individual measurement grouped by biological replicate (5 replicates per condition, 5 measurements per replicate). Horizontal black lines indicate per-replicate means. Within-sample measurements cluster tightly, demonstrating high spatial reproducibility of AFM measurements. Between-sample variance is larger, particularly in riboflavin-treated conditions, consistent with biological heterogeneity. Dotted vertical lines separate treatment groups. (D) Stiffness after low-power UV (0.3 mW/cm^2) exposure. Bars represent mean +/− SEM with individual data points overlaid. Fold-change relative to baseline (∼52 kPa, dashed line) is indicated above each bar.

### Riboflavin-UVA treatment produced concentration-dependent stiffening

UV-A exposure at 3 mW/cm² without riboflavin produced a modest increase in tissue stiffness. In PBS-treated control samples, the elastic modulus increased from 51.5 ± 2.6 kPa at baseline to 123.1 ± 2.6 kPa after UVA, corresponding to an approximately 2.4-fold increase over baseline (mean difference = 71.7 kPa, 95% CI [64.2, 79.1]; pairedt-test, t(24) = −19.84, p = 2.2 × 10⁻^16^; **Fig. 3A–B**).

At 2 mM riboflavin, the elastic modulus increased from 52.7 ± 2.8 kPa to 2,417 ± 141 kPa (mean difference = 2,364 kPa, 95% CI [2,061, 2,667]; t(24) = −16.60, p = 9.5 × 10⁻¹⁵), a 46-fold increase over baseline. At 4 mM riboflavin, stiffness reached 4,671 ± 147 kPa (mean difference = 4,618 kPa, 95% CI [4,313, 4,923]; t(24) = −31.04, p = 4.7 × 10⁻²¹), corresponding to an 89-fold increase. At the highest concentration tested, 8 mM riboflavin, stiffness rose to 8,006 ± 162 kPa (mean difference = 7,953 kPa, 95% CI [7,617, 8,288]; t(24) = −47.25, p = 1.5 × 10⁻²⁵; **Fig. 3A**), representing an approximate 150-fold increase over baseline. The highest riboflavin concentration thus produced a 65-fold greater absolute stiffness compared to UV alone (8,006 vs. 123 kPa).

Paired before-and-after analysis confirmed that stiffening occurred within individual tissue samples rather than reflecting between-sample variability (**Fig. 3B**). In contrast, PBS-treated control samples showed only limited stiffening after UVA exposure. Together, these data indicate that UVA exposure alone has a measurable but modest effect on dural nanomechanical stiffness, whereas riboflavin-UVA treatment produces a substantially larger and concentration-dependent stiffening response.

### Post-treatment stiffness differed significantly between all treatment conditions

Post-UVA elastic modulus differed significantly across treatment groups (one-way ANOVA, F(3,96) = 654.21, p < 10⁻⁶⁰). Post hoc pairwise comparisons using Tukey’s HSD test confirmed that every pair of conditions differed significantly from one another (adjusted p < 0.001 for all six comparisons; **Table 1**). Because variance differed across groups, Welch’s t-tests were performed as a sensitivity analysis and confirmed that all pairwise comparisons remained significant. Because variance differed across groups, Welch’s t-tests were performed as a sensitivity analysis and confirmed that all pairwise comparisons remained significant (W = 19.40, p < 0.0001). All pairwise comparisons remained highly significant under the unequal-variance assumption (all p < 10⁻¹⁴), confirming that the results were robust to heteroscedasticity.

**Table 1.** Pairwise post hoc comparisons of post-UV elastic modulus across treatment conditions (Tukey’s HSD).

| Group 1 | Group 2 | Mean Diff (kPa) | Tukey p (adj) | Welch p |
| --- | --- | --- | --- | --- |
| PBS Control | 2 mM RF | 2,294 | < 0.001 | < 10<sup>-14</sup> |
| PBS Control | 4 mM RF | 4,548 | < 0.001 | < 10<sup>-14</sup> |
| PBS Control | 8 mM RF | 7,883 | < 0.001 | < 10<sup>-14</sup> |
| 2 mM RF | 4 mM RF | 2,254 | < 0.001 | < 10<sup>-14</sup> |
| 2 mM RF | 8 mM RF | 5,589 | < 0.001 | < 10<sup>-14</sup> |
| 4 mM RF | 8 mM RF | 3,335 | < 0.001 | < 10<sup>-14</sup> |

*Riboflavin (RF); mean difference (Mean Diff), in post-UV elastic modulus between groups; adjusted for multiple comparisons (adj).*

Effect sizes for riboflavin conditions relative to PBS control were uniformly very large: Cohen’s d = 4.61 (95% CI [3.60, 5.62]) for 2 mM, d = 8.74 (95% CI [7.15, 10.33]) for 4 mM, and d = 13.78 (95% CI [11.39, 16.16]) for 8 mM riboflavin. These values exceed the conventional threshold for a “large” effect (d = 0.8) by an order of magnitude, indicating that the treatment effects are not only statistically significant but practically meaningful.

### Riboflavin concentration and stiffness exhibited a linear dose–response relationship

Among riboflavin-treated samples, post-UV stiffness increased linearly with riboflavin concentration (**Fig. 3A**). Linear regression on individual measurements yielded a slope of 918 ± 36 kPa per mM (95% CI [848, 988]), with R² = 0.901 (F(1, 73) = 666.3, p = 2.0 × 10⁻³⁸), indicating that riboflavin concentration explained 90% of the variance in post-UV stiffness. Regression on group means confirmed near-perfect linearity (R² = 0.994, slope = 918 kPa/mM).

A similar dose-dependent pattern was observed at the lower UV power density of 0.3 mW/cm², although absolute stiffness values were correspondingly lower at each riboflavin concentration (PBS: 101.3 ± 1.3 kPa; 2 mM: 841.6 ± 19.1 kPa; 4 mM: 1,102.7 ± 51.5 kPa; 8 mM: 1,802.0 ± 57.9 kPa; **Fig. 3A**, **Supplementary Fig. 1D**). The complete crosslinking parameter space is summarized in **Fig. 3B**, which illustrates the interaction between riboflavin concentration and UV power density. Maximum stiffening (8,006 kPa, 150-fold over baseline) was achieved at 8 mM riboflavin with 3 mW/cm² UV. At both UV power levels, the linear dose–response relationship indicates that the degree of crosslinking can be precisely titrated by adjusting riboflavin concentration, supporting the potential for controllable, predictable tissue stiffening in clinical applications.

## Discussion

This study demonstrates that riboflavin-sensitized UVA photo-crosslinking produces a dose-dependent increase in the nanomechanical stiffness of ex vivo bovine dura mater, as measured by AFM nanoindentation. The comparable baseline distributions support the interpretation that post-treatment differences were primarily treatment-associated rather than driven by pre-existing baseline variability. Treatment with 8 mM riboflavin followed by 3 mW/cm² UVA irradiation increased the elastic modulus 150-fold over baseline, from approximately 52 kPa to over 8,000 kPa. This effect was dependent on riboflavin concentration, with stiffness scaling linearly at 918 kPa per mM (R² = 0.99 on group means; **Fig. 3A**), indicating that the degree of crosslinking can be precisely titrated. To our knowledge, this is the first demonstration that riboflavin-UVA crosslinking can be applied to the dura mater, extending the clinical principle of corneal collagen crosslinking to a new target tissue of direct neurosurgical relevance.

The magnitude of mechanical stiffening observed here substantially exceeds what has been reported for corneal collagen crosslinking using the standard Dresden protocol [21]. In the cornea, riboflavin-UVA treatment using 0.1% riboflavin (approximately 0.27 mM) and 3 mW/cm² UVA for 30 minutes typically produces a 2- to 4.5-fold increase in elastic modulus [22]. The larger fold-changes observed in the present study—ranging from 46× at 2 mM to 150× at 8 mM riboflavin—likely reflect both the substantially higher riboflavin concentrations employed and fundamental differences in tissue architecture. The cornea possesses a highly ordered lamellar arrangement of type I/V collagen fibrils with precise interfibrillar spacing maintained by proteoglycans, an organization optimized for optical transparency rather than mechanical load bearing. The dura mater, by contrast, consists of densely packed, multi-directional type I collagen fibers arranged in a more disorganized, tendon-like architecture. This structural arrangement may offer a greater density of crosslinkable sites per unit volume, allowing more covalent bonds to be formed under equivalent photochemical conditions. Additionally, the dura’s greater thickness permits deeper riboflavin penetration and a larger effective volume of crosslinked tissue, whereas corneal crosslinking is typically confined to the anterior 200–300 µm of stroma [23].

A notable finding was that UVA irradiation alone, in the absence of riboflavin, produced a modest but statistically significant 2- to 3-fold increase in stiffness (**Fig. 3A–B**). This observation is consistent with known photochemical effects of UVA on collagenous tissues, where direct absorption by endogenous chromophores (such as tryptophan and tyrosine residues) can generate reactive oxygen species capable of inducing a low level of non-specific crosslinking. Importantly, this background UV effect was small compared to the riboflavin-amplified response, confirming that riboflavin is the critical mediator of the observed stiffening. The inclusion of UV-only controls in the experimental design thus served to clearly delineate the riboflavin-specific contribution from any non-specific photochemical effects.

The near-perfect linear relationship between riboflavin concentration and post-UV stiffness (R² = 0.99 on group means) is a finding of considerable translational significance. The observation of a qualitatively similar dose-response at 0.3 mW/cm² (**Fig. 3C–D, Supplementary Fig. 1D**), albeit at lower absolute stiffness values, indicates that both riboflavin concentration and UV power density can be independently modulated to achieve a target tissue stiffness. Current approaches to dural repair, including primary suture closure, dural substitutes (e.g., collagen matrices, expanded PTFE), and fibrin sealants [24], primarily act as passive barriers to CSF leakage. They do not modify the inherent mechanical properties of the native dura. In contrast, riboflavin-UVA crosslinking would actively strengthen the host tissue, representing a fundamentally different and potentially complementary strategy. Several clinical scenarios stand to benefit most directly. In patients with connective tissue disorders such as Marfan syndrome or Ehlers–Danlos syndrome, where dural ectasia and inherent tissue fragility predispose to high rates of CSF leakage [25], a method to preemptively or intraoperatively strengthen the dural sac could improve suture-holding capacity and reduce postoperative leak rates. In revision surgeries, where the dura is often scarred, thinned, and friable, crosslinking could restore mechanical competence to tissue that would otherwise be difficult to repair primarily. Following the resection of dural-invasive tumors such as meningiomas, reinforcing the native dural margins adjacent to a graft or substitute could ensure a more robust reconstruction and potentially reduce the incidence of pseudomeningocele formation. In Chiari decompression, where duraplasty is commonly performed[26], crosslinking the native dural edges could enhance the durability of the closure. Finally, crosslinking could be applied in various SIH surgical scenarios biomechanically reinforcing the edges of the duroplasty[27–29].

Our choice of AFM-based nanoindentation as the primary measurement modality was motivated by its ability to provide direct, quantitative mechanical characterization at the tissue surface with high spatial resolution. AFM has been established as a validated tool for characterizing soft biological tissue mechanics, including collagenous membranes, with high sensitivity to regional mechanical heterogeneity. The paired experimental design—measuring identical regions before and after treatment—minimized inter-sample variability and provided strong internal controls, as evidenced by the tight within-sample clustering of replicate measurements (**Supplementary Fig. 1C**) and the absence of overlap between pre- and post-UV distributions in riboflavin-treated samples (**Fig. 3B**). An important consideration is that AFM nanoindentation measures the local elastic modulus at the tissue surface under compressive loading at the nanoscale, which is conceptually distinct from the bulk tensile modulus typically reported for dura mater. Published macro-scale tensile studies report human cranial dura elastic moduli in the range of 16–130 MPa, with a mean of approximately 60–70 MPa [30,31]. Our baseline AFM measurements of approximately 52 kPa are three orders of magnitude lower, a discrepancy that is expected and well-documented across soft biological tissues when comparing nanoindentation with bulk tensile testing. AFM probes individual collagen fibril mechanics and local matrix compliance at the surface, whereas tensile testing engages the full cross-sectional architecture, including inter-laminar and interfibrillar load transfer mechanisms that contribute to bulk stiffness. These measurements are therefore complementary rather than contradictory, and the large fold-change we observe at the nanoscale provide evidence that crosslinking is modifying the fundamental building blocks of dural mechanical integrity.

Several limitations of this study must be acknowledged. First, this work was conducted entirely on ex vivo bovine dura mater. While bovine dura is a widely accepted model for neurosurgical research due to its structural and compositional similarity to human dura [32], species-specific differences in collagen fibril diameter, packing density, and extracellular matrix composition may influence the magnitude and distribution of the crosslinking effect. Validation in human dural tissue is therefore essential before any clinical extrapolation can be made. Second, the study assessed mechanical changes using AFM nanoindentation, which provides surface-level compressive modulus measurements. While these are informative, the clinical relevance of dural strengthening ultimately depends on macro-scale functional properties such as tensile strength, burst pressure, and suture pull-out resistance. Third, our study did not assess the depth of crosslinking penetration through the full dural thickness. In corneal crosslinking, the stiffening effect is known to be depth-dependent, with the anterior stroma stiffening preferentially [23]. Fourth, this study did not assess the biocompatibility or cytotoxicity of the treatment. The UVA irradiances and riboflavin concentrations used here are considerably higher than those employed in the standard corneal crosslinking protocol. While the dura mater is a relatively acellular tissue compared to the cornea, UVA-induced reactive oxygen species could potentially damage adjacent structures, including the arachnoid membrane, cortical surface, and dural vasculature. The safety threshold for UVA exposure to neural tissue has not been established, and rigorous in vivo toxicity studies in animal models are an absolute prerequisite for any clinical application. Fifth, the long-term stability of the induced crosslinks was not evaluated. Although riboflavin-UVA crosslinking in the cornea has been shown to produce durable mechanical changes lasting years [33,34], the dura mater exists in a different biomechanical and biological environment, including exposure to CSF and potential enzymatic degradation.

This study demonstrates that riboflavin-sensitized UVA crosslinking can increase the local nanomechanical stiffness of ex vivo bovine dura mater in a dose-dependent manner. The findings establish an initial proof of concept that dural tissue mechanics can be modified by photochemical collagen crosslinking. Further studies are required to determine whether these nanomechanical changes translate into improved macroscopic repair properties, including tensile strength, burst pressure, suture pull-out resistance, and CSF leak resistance. Validation in human dura, assessment of crosslinking penetration depth, and safety testing for adjacent neural and vascular tissues will be essential before clinical translation can be considered.

## Data Availability Statement

All relevant data are within the manuscript and its supporting information files. All code and data are available in the project repository (https://github.com/kevinj24fr/dura-photocrosslinking).

